# DNA replication is not a limiting factor for rapid growth of *Vibrio natriegens*

**DOI:** 10.1101/2023.05.26.541695

**Authors:** Lea Ramming, Daniel Stukenberg, María del Carmen Sánchez Olmos, Anke Becker, Daniel Schindler

## Abstract

DNA replication prior to cell division is essential for the proliferation of all cells. Bacterial chromosomes are replicated bidirectionally from a single origin of replication, with replication proceeding at about 1000 bp per second. For the best-studied model organism, *Escherichia coli*, this translates into a replication time of about 40 min for its 4.6 Mb chromosome. Nevertheless, *E. coli* can propagate by overlapping replication cycles with a maximum short doubling time of 20 min. The fastest growing bacterium known today, *Vibrio natriegens*, is able to replicate with a generation time of less than 10 min. It has a bipartite genome with chromosome sizes of 3.2 and 1.9 Mb. Is simultaneous replication from two origins a prerequisite for its rapid growth? We fused the two chromosomes of *V. natriegens* to create a strain carrying a 5.2 Mb chromosome with a single origin of replication. Compared to the wild-type, this strain showed little deviation in growth rate. This suggests that the split genome is not a prerequisite for rapid growth, and that DNA replication is not an important growth rate-limiting factor.

## Introduction

Every cell must replicate its genome prior to cell division. Canonical initiation of DNA replication in bacteria is primed at a single origin of replication (*ori*) from which the chromosome (chr) is replicated bidirectionally once per cell cycle (Reyes-Lamothe et al., 2012). The processive rate of DNA polymerase is approximately 1000 bp per second (Baker and Bell, 1998). Since DNA must be replicated prior to completing the cell cycle, replication rate can determine generation time. To overcome this bottleneck, some bacteria have evolved a system of overlapping replication cycles to increase growth rates (Cooper and Helmstetter, 1968). Notably, initiation of DNA replication still takes place only once per cell cycle but daughter cells are already born with replicating chromosomes (Skarstad et al., 1986; Fossum et al., 2007). By maintaining high *ori:ter* ratios, *Escherichia coli*, achieves doubling times of 20 min (Cooper and Helmstetter, 1968). The genome of *E. coli* is organized in a single chromosome with a size of 4.6 Mb. In contrast, the human pathogen *V. cholerae* has a bipartite genome with chromosome sizes of 3.0 Mb and 1.1 Mb, respectively (Heidelberg et al., 2000). *V. cholerae* was reported to achieve doubling times faster than *E. coli*, and the bipartite genome may be a reason for its faster growth (Couturier and Rocha, 2006). In an earlier study researchers were able to engineer the *V. cholerae* genome into a single chromosome strain and a strain with equal sized chromosomes (Val et al., 2012). Interestingly, both strains exhibit an increased doubling time in defined rich media of 26 and 34.8%, respectively. Growth rate alteration in single-chromosome strain may be explained by the extended replication time necessary to duplicate the fused chromosome from a single *ori*.

In recent years, *Vibrio natriegens* has received increased attention because of its rapid growth with reported doubling times <10 min, despite already being known for > 60 years (Payne, 1958; Eagon, 1962; Hoff et al., 2020). *Vibrio natriegens* has a bipartite genome with chromosome sizes of 3.2 Mb and 1.9 Mb, respectively. Researchers have identified a set of 587 *V. natriegens* genes required for rapid growth in rich media, identified by CRISPRi screening (Lee et al., 2019). Among those genes are ribosomal proteins, metabolic genes, and genes of the DNA polymerase. As expected, the reduction of essential proteins such as the DNA polymerase results in reduced growth rate. However, this result does not determine whether DNA replication is a rate limiting factor for *V. natriegens’* rapid growth.

To investigate the role of DNA replication on maximum cellular growth rate, we reconfigured the chromosomal architecture to require all replication to fire from a single origin. We created and characterized the *V. natriegens* strain synSC1.0 (<u>syn</u>thetic <u>s</u>ingle <u>c</u>hromosome v.<u>1.0</u>), a strain derivative of ATCC14048 with its two chromosomes fused into a singular chromosome. We prove by replication pattern analysis that the replication of the fused chromosome is initiated from a single origin of replication. We were expecting increased doubling times in synSC1.0 based on the existing reports of work with *V. cholerae* MCV1. However, our results indicate that the consequences of extended DNA replicon length in synSC1.0 are negligible for its rapid growth. The strain synSC1.0 will allow novel approaches to study chromosome biology in this rapidly growing bacterium. *V. natriegens* may be a suitable alternative to *V. cholerae*, the currently most well-studied model organism for bipartite microbial genomes, allowing to study chromosome biology without the risk of infections. Besides its application in basic research synSC1.0 may be an interesting chassis for synthetic biology and applied research, e.g. for hosting an additional synthetic chromosome.

## Results

### Construction and validation of a single chromosome V. natriegens strain

Assuming a replication speed of 1000 bp/s, the replication time for the *E. coli* genome is approx. 40 minutes, which is in line with the literature (Baker and Bell, 1998). Transferring this replication speed to the bipartite genome of *V. natriegens* would result in a replication time of 27 minutes for the larger chr1 (3.2 Mb), while in a strain with fused chromosomes (5.2 Mb) the replication time would be approx. 43 minutes, an increase of around 60%. To test if DNA replication is the rate limiting factor of *V. natriegens* rapid growth, we fused the two chromosomes by replacing the <u>d</u>eletion-induced filamentation (*dif*) site (Kuempel et al., 1991) of chr1 with the whole chr2 except for the *ori2* region. The fused chromosome possesses *oriI* for initiation of DNA replication and *dif2* for chromosome dimer resolution (*cf*. Figure 1A). The *ori2* region contains the genes for the partitioning system ParAB_2_ and the chr2 replication initiator protein RctB. The strain construction was performed utilizing our earlier published NT-CRISPR procedure (Figure 1A). In two subsequent editing steps, we integrated homologous sequences of chr1 flanking the *dif1* site upstream and downstream of the *ori2* region in chr2. Initially we planned to enforce chromosome fusion through a gRNA directing Cas9 to the *ori2* region. Surprisingly, we did not observe any inducible cell killing and, upon further inspection, we found that the chromosomes were already fused while integrating the second homologous flank. The obtained strain was termed *V. natriegens* synSC1.0 and was verified after initial Sanger sequencing of the fusion sites by pulsed-field-gel-electrophoresis (PFGE) and long-read whole-genome sequencing (Figure 1B). The PFGE shows two bands of 1.9 and 3.2 Mb for the parental strain, which are absent in synSC1.0, showing only a single band with increased size of approx. 5.2 Mb. Long-read *de novo* assembly resulted in two circular contigs for the parental strain and a single circular contig for synSC1.0 with sizes of 3.2, 1.9 and 5.2 Mb, respectively. Analysis of the chromosome fusion region in synSC1.0 revealed the absence of *dif1* and *ori2* as well as the expected chromosome fusion regions, despite a small deletion of 38 nucleotides corresponding to the ARNold (Naville et al., 2011) predicted terminator of the deleted *rctB* gene, and was considered to be negligible (Figure 1C).

**Figure 1.**
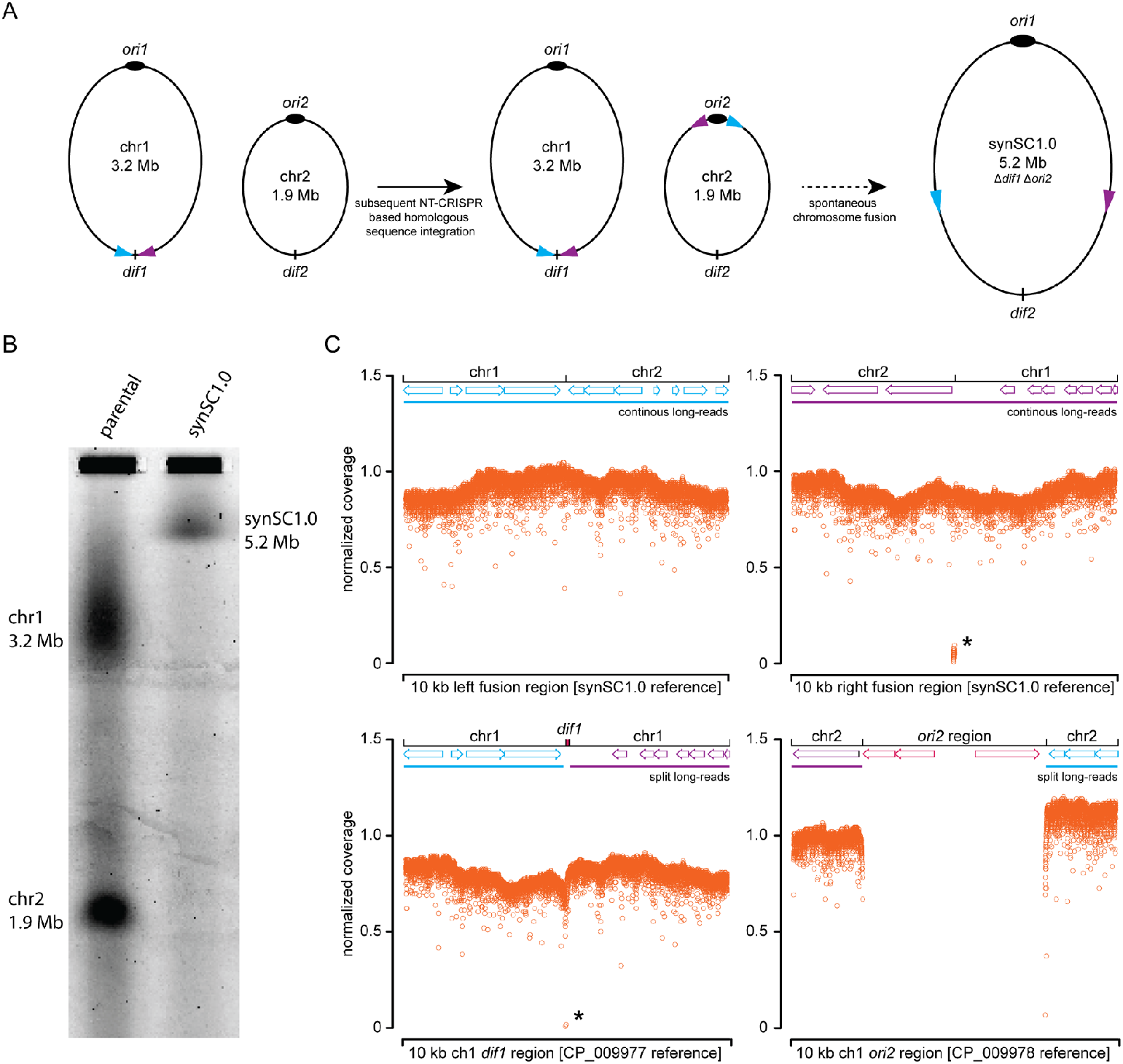
Construction and validation of the *V. natriegens* strain synSC1.0. (A) Scheme of the developed strategy for chromosome fusion and large-scale genome engineering of *V. natriegens*. NT-CRISPR is utilized to subsequently integrate homologous flanking sequences into the second chromosome in the initial step. The picked homologous sequences and their orientation are indicated by blue and purple arrows. We planned to select for the fused chromosome with deleted *dif1* and *ori2* with double strand breaks from CRISPR/Cas9. To our surprise, the chromosome fusion already occured in the initial step while integrating the second homologous region. Sizes are not to scale and differences in chromosomes sizes are due to truncation. (B) PFGE shows two bands for the parental strain with sizes of approx. 3.2 and 1.9 Mb representing chromosome 1 and 2, respectively. For synSC1.0, only a single band with an increased size of approx. 5.2 Mb is visible. The sizes were estimated by using *S. cerevisiae* and *S. pombe* samples as reference (data not shown). (C) Long-read sequencing of synSC1.0 confirms fusion of the two chromosomes and deletion of *dif1* and the *ori2* region. The two top panels show the confirmation of the left and right fusion sites visualized by the normalized coverage for a 10 kb window using the designed synSC1.0 reference. The right panel indicates the small deletion of 38 nucleotides indicated by an asterisk. The two bottom panels show the data plotted against the CP_009977 and CP_009978 references respectively to validate the deletion of *dif1* (left panel, indicated by an asterisk) and the *ori2* region (right panel). X-axis resembles a 10 kb window. Zero values are not plotted. The top of each graph contains a scheme open reading frame annotations of the genetic content in this region, while the lines indicate long-reads either spanning the whole region (continuous color, top panel) or continue at different coordinates (split color, lower panel) based on the indicated fusion sites and reference sequence. Blue and purple indicate the left and right fusion region, respectively, and deleted regions are annotated in red.

### Fusing the two chromosomes of V. natriegens results only in minor growth differences

To answer the most pressing question if the bipartite genome organization and the resulting time for DNA replication is the speed-limiting factor of *V. natriegens* rapid growth we performed comparative growth rate determination. We compared the growth of the parental strain with synSC1.0 and used *E. coli* MG1655 wild type cells as an outgroup in LBv2 media (Weinstock et al., 2016)(Figure 2A). The minimal doubling time was determined to be 12 min 11 s (+/- 34.5 s) for the parental strain and 13 min 2 s (+/- 27.1 s) for synSC1.0 under our experimental conditions. The difference in growth is 7%, which is only a fraction of the expected 60% if replication would be the speed-limiting factor for *V. natriegens* rapid growth. This difference is lower compared to the observed generation time increase for the engineered *V. cholerae* MCH1, where an increase of 26% was observed (Val et al., 2012). A detailed analysis of the growth curves indicates no drastic alteration regarding lag-phase and total biomass under the tested growth conditions (Figure 2B). In defined M9 media supplemented with 20.5 g/L NaCl and 0.4% glucose the doubling time is 22 min 46 s and 23 min 39 s for the parental and synSC1.0 strain, respectively (Figure S1). To our surprise, synSC1.0 grown under these conditions exhibited a reduced lag-phase compared to the parental strain (data not shown). Taking the increased time for DNA replication into account, we wondered if the parental strain would have an advantage under conditions causing replication stress. To test this, we assessed growth in the presence of ciprofloxacin or nalidixic acid, both gyrase inhibitors, initially in a minimum inhibitory concentration assay (MIC) (Figure 2C) and subsequently in growth assays with fine adjusted concentrations (data not shown), but could not observe growth differences between the two strains. Further, we checked if synSC1.0 possesses an increased mutation rate in fluctuation assays but could not observe significant differences in the presence of ethyl methanesulfonate (EMS) or methyl methanesulfonate (MMS), respectively. Suggesting DNA repair mechanisms are not impaired under the tested conditions (data not shown). To check whether an altered cell phenotype (e.g. elongated cell morphology) affected our optical density measurements, we performed light microscopy (Figure 2D-E). No drastic differences were observed, which is consistent with the previously described chromosome fusion of *V. cholerae* MCV1 (Val et al., 2012) but not with the drastic phenotypic alteration for a natural single chromosome isolate of *V. cholerae* observed earlier (Bruhn et al., 2018). Nevertheless, we observed individual aberrant cells in the synSC1.0 with a higher frequency compared to the parental strain (Figure 2E, Figure S2). The result may indicate an issue with chromosome segregation or chromosome dimer resolution at *dif2*. An increased number of chromosome dimers with increasing chromosome size was described previously in the study characterizing *V. cholerae* MCV1 and would match the *dif* associated filamentation phenotype (Kuempel et al., 1991; Val et al., 2012). However, taking our results together, the drastic genome rearrangement does not cause major phenotypic alterations under the tested conditions.

**Figure 2.**
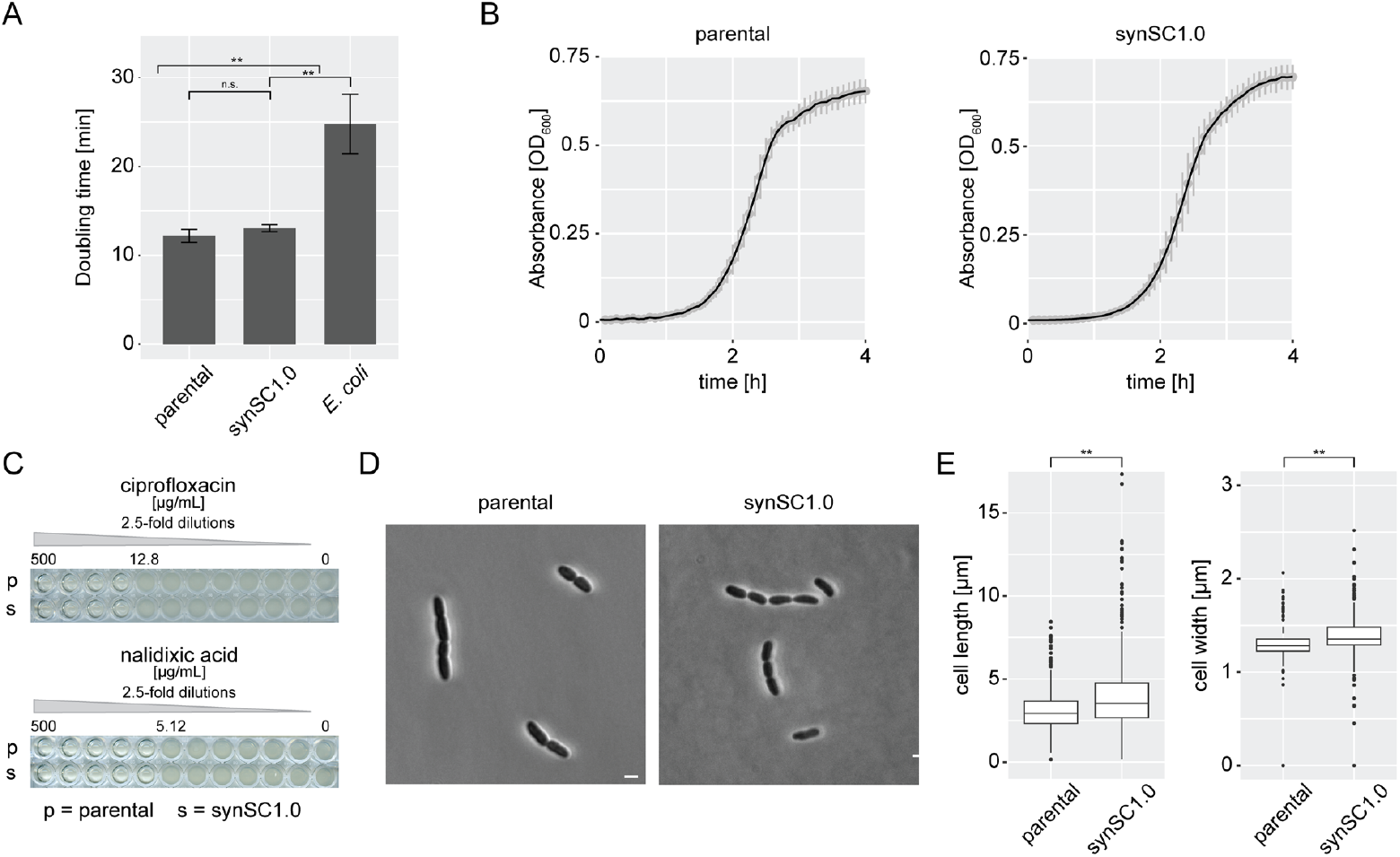
Comparative growth analysis of wild type *V. natriegens* and synSC1.0. (A) Doubling times are determined to be 12 min 11 s (+/- 34.5 s) and 13 min 2 s (+/- 27.1 s) for the parental and synSC1.0 strain, respectively. The difference in growth rate is 7%. *E. coli* was used as a control under the same conditions and doubling time was determined to be 24 min 47 s (+/- 40.1 s). Student’s t-test was applied to determine the significance; *: p< 0.01, **: p<0.001, n.s. not significant. Experiments were performed in technical triplicates with biological quadruplicates. (B) Comparison of the parental and synSC1.0 growth curve do not reveal obvious differences. The growth curves show mean of biological quadruplicates, each consisting of three technical replicates. Standard deviation is indicated by grey bars. (C) MIC determination for ciprofloxacin and nalidixic acid for parental and synSC1.0. Both substances generate DNA replication stress by inhibiting gyrase function. No differences can be observed under the tested conditions. MIC tests were performed in quadruplicates and representative examples are shown, with the highest concentration allowing growth indicated. (D) Comparison of parental strain (left panel) and synSC1.0 (right panel) cells by microscopy. There were no drastic morphological differences for the average cells. A representative image in DIC is shown for both strains. Scale bar indicates 2 μm. (E) Comparative evaluation of cell length and cell diameter for the parental and synSC1.0 strain. Values were obtained using bacstalk from four biological replicates (Hartmann et al., 2020); n = 663 (n_1_ = 32, n_2_ = 130, n_3_ = 213, n_4_ = 288) and n = 517 (n_1_ = 38, n_2_ = 103, n_3_ = 247, n_4_ = 129) for the parental and synSC1.0, respectively. The synSC1 strain shows on average a slight increased cell sizes. Student’s t-test was applied to determine the significance; *: p< 0.01, **: p<0.001, n.s. not significant.

### Replication pattern analysis indicates no differences

To prove that the *ori2* was eliminated and no cryptic *ori* is responsible for the observed growth rate we performed replication pattern analysis of synSC1.0 in comparison to the parental strain (Figure 3, Figure S3). Genomic DNA of replicating cells in early stationary phase was extracted and submitted to whole genome sequencing. Replication pattern analysis of the parental strain shows the expected pattern for chr1 and chr2 with the highest marker frequency in the regions of *ori1* and *ori2* (Figure 3A). The pattern of the data is consistent with the termination synchrony of the two chromosomes observed in *Vibrionaceae* (Kemter et al., 2018). A single peak in our replication pattern analysis of synSC1.0 verifies that the 5.2 Mb chromosome is replicated from *ori1* and no cryptic *ori* was formed (Figure 3B). Notably the stationary phase culture of synSC1.0 was not fully stationary in contrast to the parental strain and a small fraction was still replicating (Figure S3B). The *ori*:*ter* ratio was calculated based on the median reads per bin of the exponentially growing samples obtained by Repliscope for the *ori* and *ter* regions (*ori*: median of highest 100 bins, *ter*: median of lowest 200 bins). The *ori*:*ter* ratio for the parental strain was determined to be 4.15 and 2.32 for chr1 (*ori* median: 1662.5; *ter* median: 359.5) and chr2 (*ori* median: 827.5; *ter* median: 356.5), respectively. The synSC1.0 strain has an *ori*:*ter* ration of 6.65 (*ori* median: 4539.5; *ter* median: 683). The higher *ori*:*ter* ratio for synSC1.0 is expected based on the extended chromosome size resulting in a higher ratio during exponential growth; notably it is close to the addition of the ratios for chr1 and chr2 (6.47 vs 6.65). Our attempts to determine the number of replication forks using established rifampicin/cephalexin replication run-out experiments based on flow cytometry did not work (data not shown) which is consistent with reports for *V. cholerae* (Srivastava and Chattoraj, 2007; Stokke et al., 2011). The long- and short-read sequencing data of stationary phase samples were combined to perform a hybrid assembly to construct reference sequences for the parental and synSC1.0 strains with annotations based on the reference sequences of Lee *et al*. 2019. The resulting GenBank files are deposited within the BioProject PRJNA948340.

**Figure 3.**
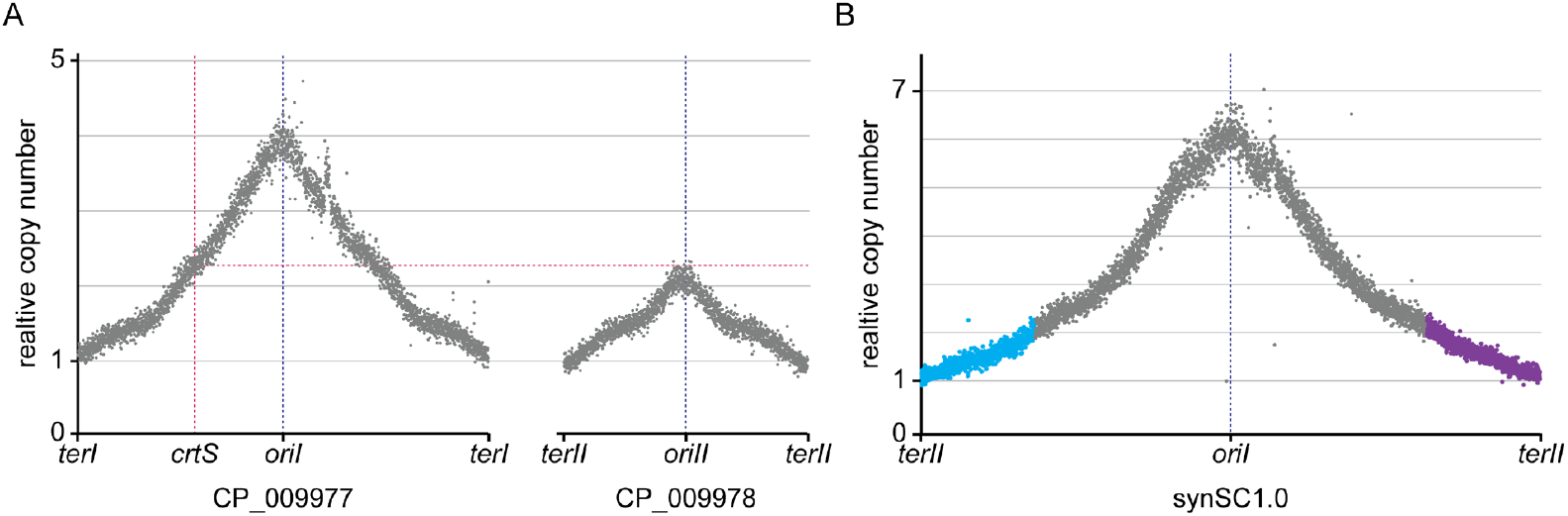
Comparative replication pattern analysis of wild type *V. natriegens* and synSC1.0. (A) and (B) show relative read numbers for 1000 bp bins for the parental strain and synSC1.0, respectively. The blue dotted lines correspond to the corresponding *ori* locations. (A) Replication pattern analysis of the wild type *V. natriegens* strain shows a single peak for each chromosome at the coordinates of the *oriI* and *oriII*. The relative copy number of *oriII* matches, as expected, to the relative copy number of location of the *crtS* site on chrI indicated by red dotted lines. (B) Replication pattern analysis of synSC1.0 shows a single peak with its maximum at the *oriI* coordinates. These results confirm the fusion of the two chromosomes, the removal of *oriII*, and the absence of alternative or cryptic *oris*. Blue and purple indicate the chr2 halves according to the fusion site color code (*cf*. Fig. 1A).

## Discussion

*V. natriegens* is the fastest growing bacterium known today and possesses a bipartite genome (Eagon, 1962; Hoff et al., 2020). We fused the two chromosomes into a single chromosome replicated from a single *ori* and were expecting a strongly reduced growth rate for *V. natriegens*. However, the growth rate only slightly deviated from that of the parental strain. This is in contrast to what was observed in *V. cholerae*, where a bigger difference was measured (Val et al., 2012). We conclude that the growth rate of the wild type *V. natriegens* is not limited by DNA replication even with a 60% increased replicon size in our study.

It seems that our engineering approach does not disrupt the described replication-associated gene dosage relevant for the fast growth in *V. natriegens*, which is in contrast to what was described to be relevant for *V. cholerae* growth rate (Couturier and Rocha, 2006). This is in line with the findings of Lee *et al*. 2019 that the content of chr2 is less relevant for *V. natriegens* rapid growth. The gene dose of chr2 encoded genes under rapid growth conditions is reduced in synSC1.0 (*cf*. Figure 3). This may result in reduced expression causing lower protein abundances. The genes encoded on chr2 could be required for niche adaptation but seem to be dispensable for growth under rich or laboratory conditions. It is tempting to speculate that the reorganized chromosome configuration of synSC1.0 lowers expression of these dispensable genes thereby liberating resources for growth related processes like ribosomes and enzymes of central metabolism, which in itself could increase growth rate. The very minor decrease in growth rate of synSC1.0 compared to the parental strain could therefore be a combination of a beneficial effect of lowering expression of non-essential genes of chr2 and the potentially detrimental effect of an increased replicon size.

The synSC1.0 strain characterized herein is a valuable resource to study chromosome biology in fast growing bacteria. synSC1.0 allows for systematic genome engineering approaches by using the *ori2* region to build synthetic, single copy chromosomes and utilize the constructs as expression platforms or to relocate and isolate genes for distinct biological functions for their in depth characterization (Messerschmidt et al., 2015; Schindler et al., 2018; Schindler, 2020). Recently, developed genetic engineering tools such as the NT-CRISPR procedure and a reusable modular cloning part collection make this organism highly accessible (Stukenberg et al., 2021; Stukenberg et al., 2022). Further, genome-scale modeling was established giving insights into *V. natriegens* metabolism and making this organism more predictable (Coppens et al., 2023). An additional advantage over the well-established *V. cholerae* model is the lack of pathogenicity allowing to conduct work in a biosafety level 1 environment, making synSC1.0 and its parental strain relevant to biotechnological and synthetic biology applications (Weinstock et al., 2016; Hoffart et al., 2017; Hoff et al., 2020; Thoma and Blombach, 2021). The halophilic nature of *V. natriegens* allows the use of sea water for its cultivation, making it a relevant emerging host in regard to a sustainable bioeconomy (Meng et al., 2022).

## Material and Methods

### Strains and growth conditions

*V. natriegens* was routinely grown in LB supplemented with v2 salts (204 mM NaCl, 4.2 mM KCl, and 23.14 mM MgCl_2_)(Weinstock et al., 2016) or in M9 media supplemented with 20.5 g/L NaCl and the indicated carbon source. Chloramphenicol was added to a final concentration of 4 μg/mL for liquid and 2 μg/mL for solid medium if applicable. Standard *E. coli* laboratory strains were used for cloning, propagation and archiving of plasmid DNA, all strains used and constructed in this study are provided in table 1. Cultures were incubated at 37 °C and at 200 rpm in case of liquid cultures if not stated otherwise. Growth comparison of *E. coli* and *V. natriegens* was performed with v2 salt containing or NaCl supplemented media for all strains.

**Table 1.** Strains used and generated in this study.

| Name | Relevant features | Parental strain | Reference |
| --- | --- | --- | --- |
| *E. coli* K12 MG1655 | K-12 F<sup>-</sup> λ<sup>-</sup> |  | (Blattner et al., 1997) |
| *E. coli* NEB Turbo | K-12 *glnV44 thi-1 Δ(lac-proAB) galE15 galK16 R(zgb-210::Tn10)Tet<sup>S</sup> endA1 fhuA2 Δ(mcrB-hsdSM)5(r<sub>K</sub><sup>-</sup>m<sub>K</sub><sup>-</sup>) F'[traD36 proAB<sup>+</sup> lacI<sup>q</sup> lacZΔM15]* |  | New England Biolabs (#C2984H) |
| *V. natriegens* Δ*dns* | Δ*dns* | *V. natriegens* ATCC14048 | (Stukenberg et al., 2021) |
| *V. natriegens*, DST026 | Δ*dns* with 3' homology flank integrated between PN96_16275 and PN96_16280 | *V. natriegens* Δ*dns* | this study |
| *V. natriegens* synSC1.0 | Δ*dns* Δ*dif1* Δ*ori2* | *V. natriegens*, DST026 | this study |

### Plasmids and oligonucleotides used in this study

Oligonucleotides were ordered and synthesized by Integrated DNA Technologies in 25 or 100 nM scale as standard desalted oligonucleotides (Tab. S1). All of the plasmids used in this study are listed in table 2 and the plasmid files of the created plasmids are provided as GenBank files with the supporting data.

**Table 2.** Plasmids used and generated in this study.

| Name | Relevant features | Parental plasmid | Reference |
| --- | --- | --- | --- |
| pST_116 + 892/893 | NT-CRISPR plasmid with gRNA for integration of 3' homology flank. Created by oligo annealing of oDS_892 and oDS_893. | pST_116_LVL2 cam | (Stukenberg et al., 2022) |
| pST_116 + 1420/1421 | NT-CRISPR plasmid with gRNA for integration of 5' homology flank. Created by oligo annealing of oDS_1420 and oDS_1421. | pST_116_LVL2 cam | (Stukenberg et al., 2022) |

### Genetic engineering of V. natriegens

tDNAs for integration of homology flanks through NT-CRISPR were generated by overlap extension PCR (Higuchi et al., 1988). Sequences of tDNAs are provided in the supporting data as GenBank files. The construction of NT-CRISPR plasmids with gRNAs targeting the integration sites were constructed as described previously through annealing of oligonucleotides (Stukenberg et al., 2022). Annealing reactions were set up by mixing 1.5 μL of each oligonucleotide (100 μM) with 5 μL T4-DNA ligase buffer (Thermo Scientific) in a total reaction volume of 50 μL. Reactions are incubated in a heat block at 95 °C for 15 min, before switching off the heat block for slowly cooling down the samples to room temperature (∼1 h). Cloning reaction with the NT-CRISPR plasmids was set up with ∼200 ng of the respective plasmid, 3 μL annealed oligonucleotides, 0.5 μL of T4-DNA Ligase (5 Weiss U/μL, Thermo Scientific) and BsaI (10 U/μL) and 1 μL T4-DNA ligase buffer in a total reaction volume of 10 μL. Reactions were run in a thermocycler with 30 cycles of 37 °C (2 min) and 16 °C (5 min), followed by a final digestion step at 37 °C for 30 min and an enzyme denaturation step at 80 °C for 10 min.

The integration of the homologous flanks were performed as described for the NT-CRISPR method (Stukenberg et al., 2022). tDNAs consisted of 3 kb homologous flanks and 3 kb insert sequence. The insert sequence is identical to the sequence upstream and downstream of *dif1* and enable fusion of chromosomes. At first, we integrated the 3’ homology flank and subsequently the 5’ homology flank. Successful integration of the 3’ homology flank was confirmed by cPCR with oligonucleotides oDS_920 and oDS_921. Integration of 5’ homology flanks and the spontaneous fusion of both chromosomes was confirmed with the primer pairs oDS_914/oDS_915 and oDS_916/oDS_917. Generation of a PCR fragment for both primer pairs indicates successful integration of 5’ homology flank without chromosome fusion, while the absence of a band for oDS_916/oDS_917 indicates chromosome fusion. Chromosome fusion was subsequently verified through Sanger sequencing (Microsynth Seqlab). Two PCR fragments spanning the junctions were generated with the primer pairs oDS_1462/oDS_1464 and with oDS_1460 and oDS_1461, each amplicon was sequenced with two reactions with primers oDS_1463/oDS_1477 and oDS_1455/oDS_1459, respectively.

### Pulsed-field-gel-electrophoresis

Plug preparation for yeast standards was performed according to described methods by Hage and Houseley (Hage and Houseley, 2013). Bacterial plug preparation was performed similar, with the following alterations: Cultures were grown overnight at 30 °C with 200 rpm. An equivalent of 1 mL OD_600_ = 5 was harvested and used for plug preparation. The concentration of low melting agarose (SeaKem LE, Lonza) for plug preparation was reduced to 0.8% and lysozyme (1 mg/mL) was used instead of lyticase. PFGE was undertaken by running samples on a 0.8% agarose gel using Pulsed Field Certified Agarose (Bio-Rad) in 1X TAE buffer at 14 °C on a Bio-Rad clamped homogeneous electric field apparatus (CHEF-DR III, Bio-Rad). 3 V/cm were used with 96 h switch time of 600 s at 120°. The resulting gel was stained with 1X SYBR Safe (ThermoFisher Scientific) and imaged using Typhoon RGB laser scanning system. The known karyotypes of *S. cerevisiae* and *S. pombe* served as size standards.

### Nanopore sequencing and data analysis

*V. natriegens* strains were cultured in 10 mL LBv2 overnight. DNA was extracted using the Monarch Genomic DNA Purification Kit (NEB) according to the manufacturer guidelines. Each sample was split in 4 purifications which were pooled subsequently again. 2 μg gDNA, corresponding to a 5-fold increase to the recommended input DNA was used as input for the library preparation using the SQK-LSK109 kit; the reason for the increase was the use of approx. 50 kb high molecular weight DNA compared to the 10 kb sized input DNA according to the protocol. The remaining procedure was performed according to the manufacturer guidelines. Each sample was sequenced on a single Flongle flow cell (FLO-FLG001 (R9.4.1)). Basecalling of raw sequencing data was performed utilizing guppy (version 6.4.6+ae70e8f; Oxford Nanopore Technologies). Basecalled raw data are deposited in BioProject PRJNA948340 individual accession IDs are provided in table S2. Initial *de novo* assembly was performed with canu (version 2.2)(Koren et al., 2017) resulting in two and one circular chromosomes for the parental and synSC1.0 strains respectively. Dot blots of *de novo* assemblies in comparison to the corresponding references based on *V. natriegens* ATCC 14048 reference sequences (CP009977 and CP009978) and the *in silico* designed single chromosome reference (deposited under BioProject PRJNA948340, *cf*. table S2) were performed using mummer (version: 3.5)(Delcher et al., 1999). Analysis of dot blots indicated duplicated segments which were later identified as assembly artifacts (data not shown) and were the reason to generate reference sequences by combining long and short-read sequencing data of this study (*cf*. section *hybrid assembly and reference construction*).

### Plate reader based growth assays

Plate reader based growth assays were adjusted to *Vibrio natriegens* based on our previously published procedure (Köbel et al., 2022; Brück et al., 2023). Briefly, *Vibrio natriegens* precultures were inoculated from a single colony and grown for 6 hours in LBv2 at 37 °C with 200 rpm. Cultures were arranged in microtiter plates and subsequently inoculated into clear, flat bottom microtiter plates (#655185, Greiner Bio-One GmbH) using a Rotor HDA+ screening robot (Singer Instruments) containing the indicated media and supplements. Plates were sealed using a PlateLoc plate sealer (Agilent) with optical clear seal. Growth was monitored in ClarioStar Plus plate readers (BMG) equipped with specific plate holders for extensive kinetics under shaking conditions. Different settings were extensively tested (data not shown) prior the following settings were identified to be the best conditions for *V. natriegens* with our setup and used throughout the study: 2 min linear shaking prior OD_600_ measurement, 800 rpm orbital shaking during idle time at 37 °C, cycle time was set to 5 min and kinetic was monitored for up to 24 hours. Raw data was exported and analyzed in R with the growthcurver package (v0.3.1)(Sprouffske and Wagner, 2016). All experiments were performed in biological quadruplicates each with technical triplicates. *E. coli* K-12 MG1655 served as an external control.

### Minimum inhibitory concentration (MIC) assay

MIC tests were performed as kinetic in ClarioStar Plus plate readers (BMG) with the settings described above for plate reader based growth assays. The only alteration was the preparation of the microtiter plate where the broth dilution method was used to determine the MIC as described previously (Köbel et al., 2022). The rationale behind this procedure was to be able to analyze growth in detail in contrast to only perform an endpoint measurement. In addition, microtiter plates were scanned at the end of the assay using an Epson Perfection V700 Photo scanner. All experiments were performed in biological quadruplicates.

### Rifampicin fluctuation assay to determine mutation frequency

Bacterial cultures were grown overnight from a single colony. 3 mL of LBv2 media without substance or test conditions (EMS [1:1,000](Sigma-Aldrich, #M0880) or MMS [1:10,000](Sigma-Aldrich, #129925)) were inoculated 1:1,000 and grown for 6 h at 37 °C with 200 rpm, respectively. 100 μL of respective dilutions were plated onto LBv2 media (10^−6^ to 10^−8^) with and without 50 μg/mL rifampicin (10^0^ to 10^−1^). The mutation frequency was determined based on CFUs, documented using an photo system (PhenoBooth, Singer Instruments). All experiments were performed in biological quadruplicates.

### Microscopic imaging and analysis

*V. natriegens* precultures were grown overnight from a single colony, inoculated 1:100 in LBv2 media and grown at 37 °C with 200 rpm for 1.5 h. 1.5 μL of exponential phase cultures were immobilized on 2% low gelling agarose (Sigma) pads containing LBv2 media and analyzed using an Axioplan 2 phase contrast microscope (Zeiss) and a Plan Neofluar 100X objective (Zeiss). Extraction of cell length and width was performed using bacstalk (Hartmann et al., 2020).

### Replication pattern analysis

Cultures for extraction of genomic DNA for replication pattern analysis were started from an overnight culture in 5 mL LBv2, (16 h, 37 °C, 200 rpm) to an OD_600_ of 0.001 in 100 mL in 1 L baffled shake flasks. Samples for exponential phase were taken after 2.5 h (OD_600_ ≈ 0.5). Samples for stationary phase were taken after 9.75 h (OD_600_ ≈ 14). A culture volume equivalent to 1 mL of OD_600_ = 2 was harvested by centrifugation for 1 min at 20,000 x g at 4 °C. Supernatant was discarded and pellet was stored at -80 °C. DNA was extracted using the Monarch Genomic DNA Purification Kit (NEB) according to the manufacturer guidelines. Library generation and short-read sequencing was performed by an external service provider with a PCR-free 150 paired-end sequencing workflow (Novogene). Replication pattern analysis was performed with Repliscope (v1.1.1)(Müller et al., 2014). All short-read sequencing raw data are deposited in BioProject PRJNA948340 and individual accession IDs are provided in table S2.

### Hybrid assembly and reference construction

Flye (version 2.9.1-b1780) was used for *de novo* assembly of long-reads (Kolmogorov et al., 2019). Resulting assemblies were corrected against the corresponding references using RagTag (version 2.1.0)(Alonge et al., 2019), respectively; *V. natriegens* ATCC 14048 (CP009977 and CP009978)(Lee et al., 2019); *V. natriegens* synSC1.0 (SAMN35394727). Polypolish (version 0.5.0) was used with standard settings to obtain polished reference genomes for both strains using short-reads from stationary phase samples (Tab. S2)(Wick and Holt, 2022). Validation and quality assessment of the assemblies was performed using Quast (version 5.2.0)(Gurevich et al., 2013). The origin of replication of each reference were set to nucleotide +1, resulting references are deposited within the BioProject PRJNA948340 (Tab. S2).

## Supporting information

- Supporting information (Figures S1-S3 and tables S1-S2)
- Supporting data (GenBank files of plasmids and tDNAs)

## Data availability

The data underlying this study are available in the published article and its online supplementary material. Sequencing raw reads and constructed reference sequences are deposited under BioProject PRJNA948340. All material created within this study is available from the corresponding author upon request.

## Acknowledgments

We thank the Becker and Schindler research groups and the MaxGENESYS biofoundry team for fruitful and inspiring discussions. We thank Tania Köbel for technical support throughout the study, Adán Andrés Ramírez-Rojas for performing the Nanopore sequencing and Timon Alexander Lindeboom for help with the pulsed-field-gel-electrophoresis. We are thankful to Christoph Klaus Spahn for support with microscopic imaging and Scott Scholz for carefully reading the manuscript.

## Contributions

DSc and DSt conceived, planned and designed the study with input of AB. LR, DSt and DSc performed experiments and analyzed the data with support of MCSO. DSc and DSt wrote the manuscript with input of all authors. All authors approved the final manuscript.

## References

Alonge, M., Soyk, S., Ramakrishnan, S., Wang, X., Goodwin, S., Sedlazeck, F.J., et al. (2019). RaGOO: Fast and accurate reference-guided scaffolding of draft genomes. Genome Biol 20(1), 224. doi: 10.1186/s13059-019-1829-6.

Baker, T.A., and Bell, S.P. (1998). Polymerases and the replisome: machines within machines. Cell 92(3), 295–305. doi: 10.1016/s0092-8674(00)80923-x.

Blattner, F.R., Plunkett, G., 3rd, Bloch, C.A., Perna, N.T., Burland, V., Riley, M., et al. (1997). The complete genome sequence of Escherichia coli K-12. Science 277(5331), 1453–1462. doi: 10.1126/science.277.5331.1453.

Brück, M., Berghoff, B.A., and Schindler, D. (2023). In silico design, in vitro construction and in vivo application of synthetic small regulatory RNAs in bacteria. arXiv. doi: 10.48550/arXiv.2304.14932.

Bruhn, M., Schindler, D., Kemter, F.S., Wiley, M.R., Chase, K., Koroleva, G.I., et al. (2018). Functionality of two origins of replication in Vibrio cholerae strains with a single chromosome. Front Microbiol 9, 2932. doi: 10.3389/fmicb.2018.02932.

Cooper, S., and Helmstetter, C.E. (1968). Chromosome replication and the division cycle of Escherichia coli B/r. J Mol Biol 31(3), 519–540. doi: 10.1016/0022-2836(68)90425-7.

Coppens, L., Tschirhart, T., Leary, D.H., Colston, S.M., Compton, J.R., Hervey, W.J.t., et al. (2023). Vibrio natriegens genome-scale modeling reveals insights into halophilic adaptations and resource allocation. Mol Syst Biol 19(4), e10523. doi: 10.15252/msb.202110523.

Couturier, E., and Rocha, E.P. (2006). Replication-associated gene dosage effects shape the genomes of fast-growing bacteria but only for transcription and translation genes. Mol Microbiol 59(5), 1506–1518. doi: 10.1111/j.1365-2958.2006.05046.x.

Delcher, A.L., Kasif, S., Fleischmann, R.D., Peterson, J., White, O., and Salzberg, S.L. (1999). Alignment of whole genomes. Nucleic Acids Res 27(11), 2369–2376. doi: 10.1093/nar/27.11.2369.

Eagon, R.G. (1962). Pseudomonas natriegens, a marine bacterium with a generation time of less than 10 minutes. J Bacteriol 83(4), 736–737. doi: 10.1128/jb.83.4.736-737.1962.

Fossum, S., Crooke, E., and Skarstad, K. (2007). Organization of sister origins and replisomes during multifork DNA replication in Escherichia coli. EMBO J 26(21), 4514–4522. doi: 10.1038/sj.emboj.7601871.

Gurevich, A., Saveliev, V., Vyahhi, N., and Tesler, G. (2013). QUAST: Quality assessment tool for genome assemblies. Bioinformatics 29(8), 1072–1075. doi: 10.1093/bioinformatics/btt086.

Hage, A.E., and Houseley, J. (2013). Resolution of budding yeast chromosomes using pulsed-field gel electrophoresis. Methods Mol Biol 1054, 195–207. doi: 10.1007/978-1-62703-565-1_13.

Hartmann, R., van Teeseling, M.C.F., Thanbichler, M., and Drescher, K. (2020). BacStalk: A comprehensive and interactive image analysis software tool for bacterial cell biology. Mol Microbiol 114(1), 140–150. doi: 10.1111/mmi.14501.

Heidelberg, J.F., Eisen, J.A., Nelson, W.C., Clayton, R.A., Gwinn, M.L., Dodson, R.J., et al. (2000). DNA sequence of both chromosomes of the cholera pathogen Vibrio cholerae. Nature 406(6795), 477–483. doi: 10.1038/35020000.

Higuchi, R., Krummel, B., and Saiki, R.K. (1988). A general method of in vitro preparation and specific mutagenesis of DNA fragments: Study of protein and DNA interactions. Nucleic Acids Res 16(15), 7351–7367. doi: 10.1093/nar/16.15.7351.

Hoff, J., Daniel, B., Stukenberg, D., Thuronyi, B.W., Waldminghaus, T., and Fritz, G. (2020). Vibrio natriegens: An ultrafast-growing marine bacterium as emerging synthetic biology chassis. Environ Microbiol 22(10), 4394–4408. doi: 10.1111/1462-2920.15128.

Hoffart, E., Grenz, S., Lange, J., Nitschel, R., Muller, F., Schwentner, A., et al. (2017). High substrate uptake rates empower Vibrio natriegens as production host for industrial biotechnology. Appl Environ Microbiol 83(22). doi: 10.1128/AEM.01614-17.

Kemter, F.S., Messerschmidt, S.J., Schallopp, N., Sobetzko, P., Lang, E., Bunk, B., et al. (2018). Synchronous termination of replication of the two chromosomes is an evolutionary selected feature in Vibrionaceae. PLoS Genet 14(3), e1007251. doi: 10.1371/journal.pgen.1007251.

Köbel, T.S., Melo Palhares, R., Fromm, C., Szymanski, W., Angelidou, G., Glatter, T., et al. (2022). An easy-to-use plasmid toolset for efficient generation and benchmarking of synthetic small RNAs in bacteria. ACS Synth Biol 11(9), 2989–3003. doi: 10.1021/acssynbio.2c00164.

Kolmogorov, M., Yuan, J., Lin, Y., and Pevzner, P.A. (2019). Assembly of long, error-prone reads using repeat graphs. Nat Biotechnol 37(5), 540–546. doi: 10.1038/s41587-019-0072-8.

Koren, S., Walenz, B.P., Berlin, K., Miller, J.R., Bergman, N.H., and Phillippy, A.M. (2017). Canu: Scalable and accurate long-read assembly via adaptive k-mer weighting and repeat separation. Genome Res 27(5), 722–736. doi: 10.1101/gr.215087.116.

Kuempel, P.L., Henson, J.M., Dircks, L., Tecklenburg, M., and Lim, D.F. (1991). dif, a recA-independent recombination site in the terminus region of the chromosome of Escherichia coli. New Biol 3(8), 799–811.

Lee, H.H., Ostrov, N., Wong, B.G., Gold, M.A., Khalil, A.S., and Church, G.M. (2019). Functional genomics of the rapidly replicating bacterium Vibrio natriegens by CRISPRi. Nat Microbiol 4(7), 1105–1113. doi: 10.1038/s41564-019-0423-8.

Meng, W., Zhang, Y., Ma, L., Lu, C., Xu, P., Ma, C., et al. (2022). Non-sterilized fermentation of 2,3-butanediol with seawater by metabolic engineered fast-growing Vibrio natriegens. Front Bioeng Biotechnol 10, 955097. doi: 10.3389/fbioe.2022.955097.

Messerschmidt, S.J., Kemter, F.S., Schindler, D., and Waldminghaus, T. (2015). Synthetic secondary chromosomes in Escherichia coli based on the replication origin of chromosome II in Vibrio cholerae. Biotechnol J 10(2), 302–314. doi: 10.1002/biot.201400031.

Müller, C.A., Hawkins, M., Retkute, R., Malla, S., Wilson, R., Blythe, M.J., et al. (2014). The dynamics of genome replication using deep sequencing. Nucleic Acids Res 42(1), e3. doi: 10.1093/nar/gkt878.

Naville, M., Ghuillot-Gaudeffroy, A., Marchais, A., and Gautheret, D. (2011). ARNold: A web tool for the prediction of Rho-independent transcription terminators. RNA Biol 8(1), 11–13. doi: 10.4161/rna.8.1.13346.

Payne, W.J. (1958). Studies on bacterial utilization of uronic acids. III. Induction of oxidative enzymes in a marine isolate. J Bacteriol 76(3), 301–307. doi: 10.1128/jb.76.3.301-307.1958.

Reyes-Lamothe, R., Nicolas, E., and Sherratt, D.J. (2012). Chromosome replication and segregation in bacteria. Annu Rev Genet 46, 121–143. doi: 10.1146/annurev-genet-110711-155421.

Schindler, D. (2020). Genetic engineering and synthetic genomics in yeast to understand life and boost biotechnology. Bioengineering (Basel) 7(4). doi: 10.3390/bioengineering7040137.

Schindler, D., Dai, J., and Cai, Y. (2018). Synthetic genomics: A new venture to dissect genome fundamentals and engineer new functions. Curr Opin Chem Biol 46, 56–62. doi: 10.1016/j.cbpa.2018.04.002.

Skarstad, K., Boye, E., and Steen, H.B. (1986). Timing of initiation of chromosome replication in individual Escherichia coli cells. EMBO J 5(7), 1711–1717. doi: 10.1002/j.1460-2075.1986.tb04415.x.

Sprouffske, K., and Wagner, A. (2016). Growthcurver: An R package for obtaining interpretable metrics from microbial growth curves. BMC Bioinformatics 17, 172. doi: 10.1186/s12859-016-1016-7.

Srivastava, P., and Chattoraj, D.K. (2007). Selective chromosome amplification in Vibrio cholerae. Mol Microbiol 66(4), 1016–1028. doi: 10.1111/j.1365-2958.2007.05973.x.

Stokke, C., Waldminghaus, T., and Skarstad, K. (2011). Replication patterns and organization of replication forks in Vibrio cholerae. Microbiology (Reading) 157(Pt 3), 695-708. doi: 10.1099/mic.0.045112-0.

Stukenberg, D., Hensel, T., Hoff, J., Daniel, B., Inckemann, R., Tedeschi, J.N., et al. (2021). The Marburg Collection: A Golden Gate DNA assembly framework for synthetic biology applications in Vibrio natriegens. ACS Synth Biol 10(8), 1904–1919. doi: 10.1021/acssynbio.1c00126.

Stukenberg, D., Hoff, J., Faber, A., and Becker, A. (2022). NT-CRISPR, combining natural transformation and CRISPR-Cas9 counterselection for markerless and scarless genome editing in Vibrio natriegens. Commun Biol 5(1), 265. doi: 10.1038/s42003-022-03150-0.

Thoma, F., and Blombach, B. (2021). Metabolic engineering of Vibrio natriegens. Essays Biochem 65(2), 381–392. doi: 10.1042/EBC20200135.

Val, M.E., Skovgaard, O., Ducos-Galand, M., Bland, M.J., and Mazel, D. (2012). Genome engineering in Vibrio cholerae: A feasible approach to address biological issues. PLoS Genet 8(1), e1002472. doi: 10.1371/journal.pgen.1002472.

Weinstock, M.T., Hesek, E.D., Wilson, C.M., and Gibson, D.G. (2016). Vibrio natriegens as a fast-growing host for molecular biology. Nat Methods 13(10), 849–851. doi: 10.1038/nmeth.3970.

Wick, R.R., and Holt, K.E. (2022). Polypolish: Short-read polishing of long-read bacterial genome assemblies. PLoS Comput Biol 18(1), e1009802. doi: 10.1371/journal.pcbi.1009802.

