## Supporting Information for "DNA replication is not a limiting factor for rapid growth of *Vibrio natriegens*"


This file contains:

Figure S1-S3

Table S1-S2

Supporting References

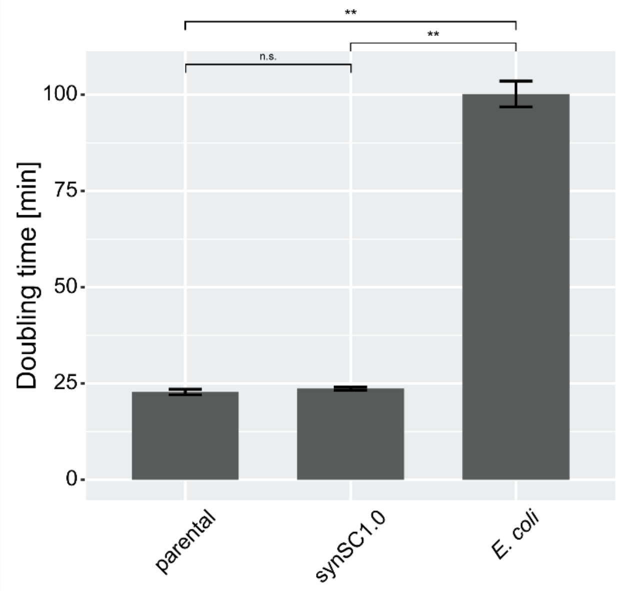


**Figure S1 | Comparative growth assay for *V. natriegens* and synSC1.0 in M9 media supplemented with 20.5 g/L NaCl and 0.4% glucose.** Doubling times are determined to be 22 min 46 s (+/- 43.8 s) and 23 min 39 s (+/- 24.6 s) for the parental and synSC1.0 strain, respectively. The difference in growth rate is 4.1%. *E. coli* was used as a control under the same conditions and doubling time was determined to be 1 h 40 min 12 s (+/- 3 min 20.8 s). However, the conditions presumably cause high salt stress for *E. coli*. Student's t-test was applied to determine the significance; *: $p < 0.01$, **: $p<0.001$, n.s. not significant. Experiments were performed biological quadruplicates each with technical triplicates.

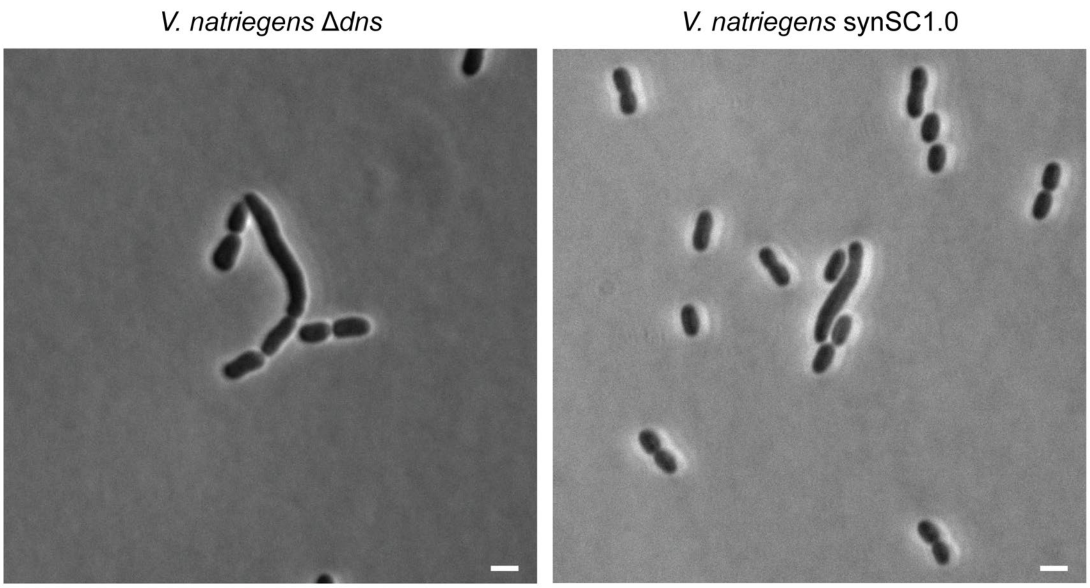


**Figure S2 | Elongated cell phenotype observed for *Vibrio natriegens* under rapid growth conditions.** Elongated cells were sporadic observed in both the parental and synSC1.0 strain. Representative image for the parental (left panel) and synSC1.0 (right panel) are shown. The observed phenotype was more frequent for synSC1.0 (*cf*. Figure 2F). It may be the result of reduced chromosome dimer resolution fidelity previously observed in *V. cholerae* (Val et al., 2012). More severe phenotypes affecting the whole cell population were described earlier for a natural isolate with fused chromosomes of *V. cholerae* but are not observed for synSC1.0 (Bruhn et al., 2018). Scale bar 2 µm.

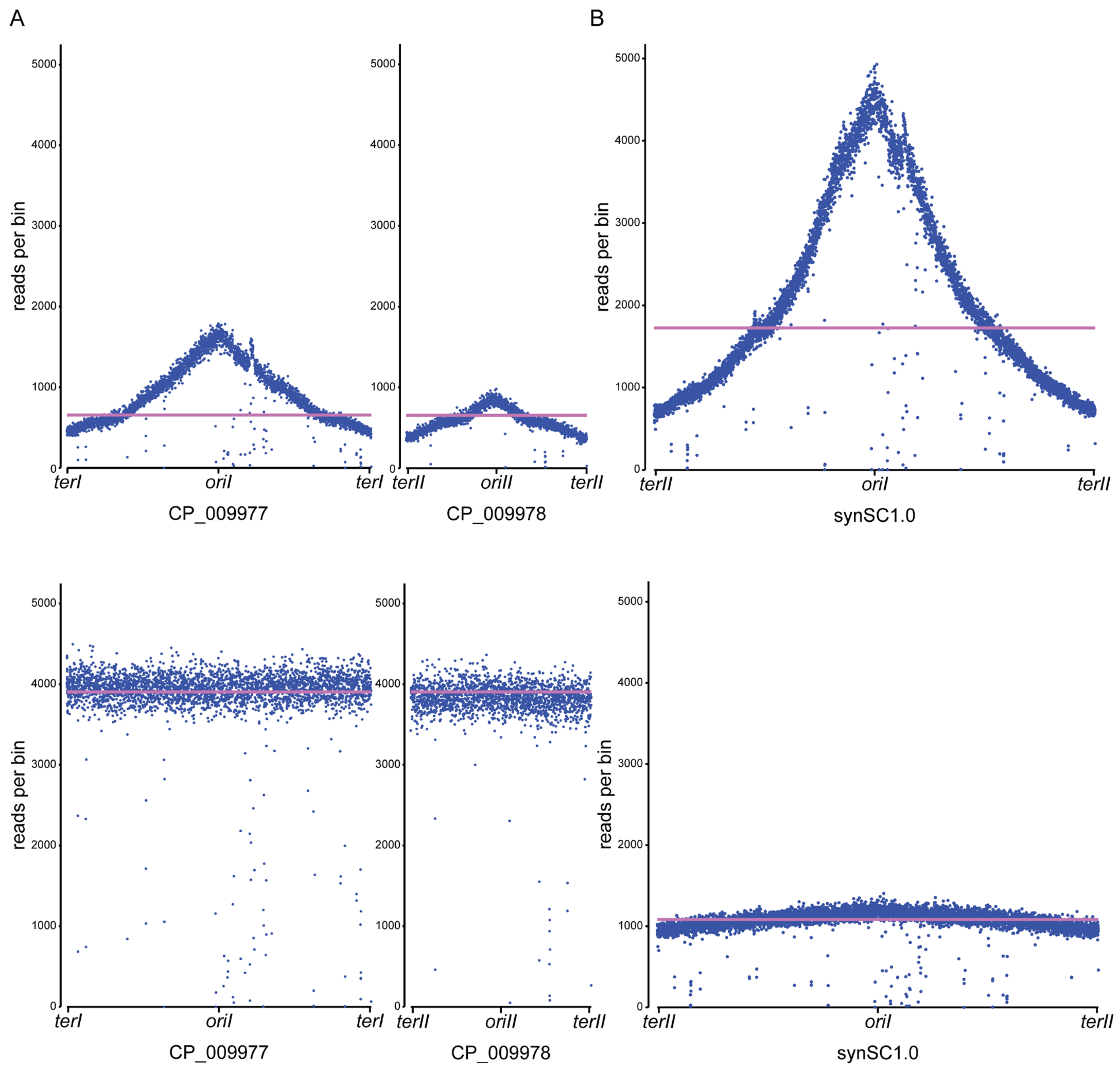


**Figure S3 | Coverage plots of samples for replication pattern analysis based on bin read count for each sample.** (A) Overall read abundances per bin (1000 bp) for *V. natriegens* parental strain in exponential growth phase (top panel) and early stationary phase (bottom panel) sequencing samples. Exponential and stationary phase samples show the expected read distribution. The magenta line indicates the sample wide median read per bin distribution. (B) Overall read abundances per bin (1000 bp) for *V. natriegens* synSC1.0 in exponential growth phase (top panel) and early stationary phase (bottom panel) sequencing samples. The stationary phase sample for synSC1.0 contained still cells undergoing DNA replication. The magenta line indicates the sample wide median read per bin distribution. All plots were generated utilizing Repliscope (Müller et al., 2014).

**Table S1 | Oligonucleotides used in this study.**

| ID | Sequence (5´-> 3´) | Purpose |
|---|---|---|
| oDS_892 | GTCCCTATCTATTAATCATCAGAA | Forward oligo for construction of gRNA. Integration of 3’ homology flank. Targeting intergenic region between PN96_16275 and PN96_16280. |
| oDS_893 | AAACTTCTGATGATTAATAGATAG | Reverse oligo for construction of gRNA. Integration of 3’ homology flank. Targeting intergenic region between PN96_16275 and PN96_16280. |
| oDS_1420 | GTCCGATCGGCAAGCAAAAACAAC | Forward oligo for construction of gRNA. Integration of 5’ homology flank. Targeting intergenic region between PN96_16260 and PN96_16265. |
| oDS_1421 | AAACGTTGTTTTTGCTTGCCGATC | Reverse oligo for construction of gRNA. Integration of 5’ homology flank. Targeting intergenic region between PN96_16260 and PN96_16265. |
| oDS_904 | GTTACAGCGTCACACATTTACATTG | Construction of tDNA for integration of 3’ homology flank. Forward primer of upstream fragment. |
| oDS_905 | TTTTGAGTCAACTTTAAATCGCCATTTCATCAAGAGC | Construction of tDNA for integration of 3’ homology flank. Reverse primer of upstream fragment. |
| oDS_906 | GCGTTATGCTTTGAATAGACTTCGCGAGTTTCTG | Construction of tDNA for integration of 3’ homology flank. Forward primer of downstream fragment. |
| oDS_907 | GAAAGTTGACAAAAACATTAACCATTGAAG | Construction of tDNA for integration of 3’ homology flank. Reverse primer of downstream fragment. |
| oDS_924 | ATGGCGATTTAAAGTTGACTCAAAAGAACCCAGCATACG | Construction of tDNA for integration of 3’ homology flank. Forward primer of insert fragment. |
| oDS_925 | GCGAAGTCTATTCAAAGCATAACGCTACGCCAAATCTTTACTC | Construction of tDNA for integration of 3’ homology flank. Reverse primer of insert fragment. |
| oDS_896 | TGACTTCATCACCAACATTGACAG | Construction of tDNA for integration of 5’ homology flank. Forward primer of upstream fragment. |
| oDS_1427 | CTTTTAACCCTTTGTTGTTTTTGCTTGCCGATCAG | Construction of tDNA for integration of 5’ homology flank. Reverse primer of upstream fragment. |
| oDS_1430 | TATAGTAGACTGTGGCATAAAAAAACCGCCCGAAG | Construction of tDNA for integration of 5’ homology flank. Forward primer of downstream fragment. |
| oDS_899 | ACCATGGATTTTATCGCCTCCTTG | Construction of tDNA for integration of 3’ homology flank. Reverse primer of downstream fragment. |
| oDS_1428 | AGCAAAAACAACAAAGGGTTAAAAGATTAACAGTGCTG | Construction of tDNA for integration of 3’ homology flank. Forward primer of upstream fragment. |
| oDS_1429 | TTTTTTTATGCCACAGTCTACTATAACCCGTATGG | Construction of tDNA for integration of 3’ homology flank. Reverse primer of insert fragment. |
| oDS_920 | CTTTCGCCCTCTAATAGAGC | Primer for cPCR. Confirming integration of 3’ homology flank. Binds in 3’ homology flank. |

| ID | Sequence (5´-> 3´) | Purpose |
|---|---|---|
| oDS_921 | GGTATCCAGCAAAGTCTGTC | Primer for cPCR. Confirming integration of 3' homology flank. Binds in chr2 sequence downstream of integration site. |
| oDS_914 | GGGATCCCTTTAGTAGCAGG | Primer for cPCR. Confirming integration of 5' homology flank. Binds in chr2 sequence upstream of integration site. |
| oDS_915 | CCAGAGCGAAGCATGAATCC | Primer for cPCR. Confirming integration of 5' homology flank. Binds in 5' homology flank. |
| oDS_916 | GGATTCATGCTTCGCTCTGG | Primer for cPCR. Confirming integration of 5' homology flank. |
| oDS_917 | CATGACATTTGTACAGGTTATCC | Primer for cPCR. Confirming integration of 5' homology flank. Binds in *oriII* region on chr2. |
| oDS_1462 | ACCCGGAAAGATGATCAAGG | Forward primer to generate PCR fragment for Sanger sequencing to confirm chromosome fusion. |
| oDS_1464 | AATCAGGTGAATTTATTAATGTAAAGGAC | Forward primer to generate PCR fragment for Sanger sequencing to confirm chromosome fusion. |
| oDS_1463 | CCAAGCAAAGTTATTAAGCTCAGC | Sequencing primer to confirm chromosome fusion. |
| oDS_1477 | CATTATGTTTAATGAAAGCCTCAAACC | Sequencing primer to confirm chromosome fusion. |
| oDS_1460 | GGGTGCCTTGCGATATAGC | Forward primer to generate PCR fragment for Sanger sequencing to confirm chromosome fusion. |
| oDS_1461 | ATTCACCGACACAAAACCAACG | Reverse primer to generate PCR fragment for Sanger sequencing to confirm chromosome fusion. |
| oDS_1455 | GAACGAAGTGATAAGTTCGTTTTGC | Sequencing primer to confirm chromosome fusion. |
| oDS_1459 | AAGTAGGGTACTTGGAACTTTTCC | Sequencing primer to confirm chromosome fusion. |

**Table S2 | Sequencing data deposited under BioProject PRJNA948340.**

| IDs | Type of data | Platform | Strain | Additional information |
|---|---|---|---|---|
| SAMN35035422 | Raw reads | Illumina (150 PE) | *V. natriegens* Δ*dns* | exponential growing sample |
| SAMN35035423 | Raw reads | Illumina (150 PE) | *V. natriegens* Δ*dns* | stationary phase sample |
| SAMN35035424 | Raw reads | Illumina (150 PE) | *V. natriegens* synSC1.0 | exponential growing sample |
| SAMN35035425 | Raw reads | Illumina (150 PE) | *V. natriegens* synSC1.0 | stationary phase sample |
| SAMN35035426 | Raw reads | Nanopore (MinION) | *V. natriegens* Δ*dns* | stationary phase sample |
| SAMN35035427 | Raw reads | Nanopore (MinION) | *V. natriegens* synSC1.0 | stationary phase sample |
| SAMN35394727 | Reference | NA | Designed *V. natriegens* synSC1.0 | designed based on CP009977 and CP009978 (Lee et al., 2019) |
| SAMN35394728 | Reference | NA | *V. natriegens* synSC1.0 | sequencing validated reference |
| SAMN35394729 | Reference | NA | *V. natriegens* Δ*dns* | sequencing validated reference |

### Supporting References


Bruhn, M., Schindler, D., Kemter, F.S., Wiley, M.R., Chase, K., Koroleva, G.I., et al. (2018). Functionality of two origins of replication in *Vibrio cholerae* strains with a single chromosome. *Front Microbiol* 9**,** 2932. doi: 10.3389/fmicb.2018.02932.

Lee, H.H., Ostrov, N., Wong, B.G., Gold, M.A., Khalil, A.S., and Church, G.M. (2019). Functional genomics of the rapidly replicating bacterium *Vibrio natriegens* by CRISPRi. *Nat Microbiol* 4(7)**,** 1105-1113. doi: 10.1038/s41564-019-0423-8.

Müller, C.A., Hawkins, M., Retkute, R., Malla, S., Wilson, R., Blythe, M.J., et al. (2014). The dynamics of genome replication using deep sequencing. *Nucleic Acids Res* 42(1)**,** e3. doi: 10.1093/nar/gkt878.

Val, M.E., Skovgaard, O., Ducos-Galand, M., Bland, M.J., and Mazel, D. (2012). Genome engineering in *Vibrio cholerae*: A feasible approach to address biological issues. *PLoS Genet* 8(1)**,** e1002472. doi: 10.1371/journal.pgen.1002472.
